# Inexpensive High-Throughput Multiplexed Cytokine Detection for Tuberculosis Diagnostics Using Amplified Enzymatic Metallization

**DOI:** 10.64898/2026.01.27.700981

**Authors:** Kanz Elkhiyari, Balazs Kaszala, Asma Hashim, Yuvraj Rallapalli, Jeremiah Khayumbi, Joshua Ongalo, Joan Tonui, Cheryl L. Day, Aniruddh Sarkar

## Abstract

Lack of accurate yet inexpensive diagnostics remains a critical bottleneck in the control and elimination of tuberculosis (TB). Interferon-gamma release assay (IGRA), which involve measurement of the release of the cytokine, interferon-gamma (IFN-γ), in blood samples stimulated with *Mycobacterium tuberculosis* antigens, is used to detect latent TB infection (LTBI). Use of IGRA in resource-poor settings, in which TB is endemic, is however hindered by the need for specialized equipment for sensitive detection of low amounts of cytokine released. Additionally, recent research has shown the advantage of multiplexed detection of other non-IFN-γ cytokines in improving diagnostic accuracy. However, this requires even more expensive instrumentation and there are no inexpensive or point-of-care compatible multiplexed cytokine detection tools available yet. Here we develop and demonstrate a low-cost, high-throughput, multiplexed cytokine detection platform based on amplified enzymatic silver metallization on a plastic substrate. The assay is performed in microwells formed on a commonly available plastic petri dish and the dry readout of the deposited silver is obtained using a cellphone camera, thus significantly reducing overall cost and complexity of multiplexed cytokine detection. We demonstrate the ability to measure clinically relevant sub-picomolar levels of multiple cytokines, including IFN-γ, interleukin-2 (IL-2), and tumor necrosis factor alpha (TNF-α) from a small volume (<5µL) of the same blood sample used in an IGRA. Furthermore, we demonstrate the use of this assay to distinguish IGRA+ and IGRA-participant samples from a TB endemic setting.

---

Tuberculosis (TB) caused by *Mycobacterium tuberculosis* (Mtb), affects nearly 2 billion people, of which 10 million develop active TB disease. Though TB occurs worldwide, the majority of new cases occur in thirty high TB burden countries mainly in Southeast Asia and Africa^1^. Effective TB treatment exists; however, the persistently high prevalence can be partially attributed to a large diagnostic gap. TB diagnosis is further complicated by the heterogeneous disease spectrum^2^. Pathogen detection-based methods exist for screening active TB cases, but 90-95% of individuals remain asymptomatic and are considered to have latent Mtb infection (LTBI), in which Mtb is difficult to detect due to low abundance^1, 3^. ∼10% of individuals with LTBI will progress to TB, which can be prevented with antibiotic treatment if diagnosed early.

While there are no gold standard tests for diagnosing LTBI, commonly used tests are the tuberculin skin test (TST) and the interferon-gamma release assay (IGRA), both of which are cellular immunity-based approaches. The TST is low-cost and point-of-care compatible but it suffers from poor sensitivity and specificity^4^. It is prone to false positives in individuals who have received the BCG (Bacille Calmette-Guerin) vaccination, which includes much of the population in high TB burden countries. Additionally, this method requires two visits to healthcare facilities, which may not be ideal in high-burden settings. Alternatively, IGRA is performed using a blood sample. Whole blood or isolated T-cells are stimulated with a mixture of Mtb-specific antigens: early secretory antigenic target 6 (ESAT-6) and culture filtrate protein (CFP-10)^5^. Secreted interferon-gamma (IFN-γ) in response to these antigens is measured using either an enzyme-linked immunosorbent assay (ELISA) or an enzyme-linked immunosorbent spot (ELISPOT) assay. This method is sensitive and specific, but its use is hindered by its high equipment cost and need for specialized laboratories and trained technicians^6^.

Current inexpensive, point-of-care (POC) assays such commercial lateral flow assays (LFAs) often focus on humoral immunity (i.e. antibody titer). Antibody based tests have till date failed to provide sufficient sensitivity and specificity for TB diagnosis^7^. Although newer antibody Fc-based biomarker discovery approaches have shown promise in TB and other diseases, these await further confirmation in larger cohorts and translation to point-of-care detection^8-10^. Cellular immunity-based approaches focus on the production and detection of one or more cytokines. There are limited commercially available point-of-care (POC) cytokine detection tests for infectious diseases in general. While antibodies are abundant in serum (∼µg/mL), cytokines are present in significantly smaller quantities (∼pg/mL)^11^. Though there are a few lateral flow assays (LFAs) for cytokine detection, they typically have high LODs (ng/mL), limiting their clinical and diagnostic value^12, 13^. There have been recent advancements in the POC cytokine detection using microfluidic approaches including fluorescence, chemiluminescence, and impedance readouts^14-18^.

In addition to the need for inexpensive POC cytokine detection, there is also a need for multiplexed cytokine detection for better prediction of disease state. While, thus far, the commercially available blood test for TB, or the IGRA, has been limited to IFN-γ measurement, studies have shown the value of other cytokines, such as tumor necrosis factor-alpha (TNF-α) and interleukin-2 (IL-2) as biomarkers for LTBI and for a more comprehensive diagnosis of TB. Examples include potentially differentiating between TB and LTBI, profiling drug-resistant TB, and predicting progression from latent to active TB disease^19-21^. Though there are lab-based multiplexable assays such as bead-based assays, such a multiplexed measurement of secreted cytokines currently can only be performed in a research context and is not available as a POC test.

Here, we set out to develop a low-cost, high-throughput, multiplexable cytokine detection platform for TB diagnosis. Additionally, our goal was to align this with the sample type requirement of the commonly used IGRA, while significantly reducing the resources required to run a multiplexed cytokine assay. Thus, we sought to bypass the need for expensive instrumentation such as flow cytometers and plate readers as well as to reduce the sample volume with an eventual goal of being able to perform such an assay from a drop of blood. We have previously reported multiplexed antibody-based biomarker assays using enzymatic metallization for electronic and optical detection, including the MO-BEAM (Multiplexed Optical Bioassay using Enzymatic Metallization) glass slide-based platform and the BEAD-EM (Bead-based Electronic Bio-Assay Detection using Enzymatic Metallization) bead-based platform^22-25^. Here we developed a multiplexable, signal amplification cascade on a high-binding inexpensive plastic substrate which enhances the sensitivity of enzymatic metallization-based immunoassays by 100-1000-fold, enabling detection of cytokine concentrations in the picogram per milliliter (or sub-picomolar) range. We term it as **<u>e</u>**nhanced MO-BEAM or eMO-BEAM. We report here the development of the eMO-BEAM platform and its characterization, first using model binding assays as well as assays for detecting recombinant cytokines. Detection of stimulated cell-secreted cytokines from cultured primary human immune cells is then demonstrated. Finally, detection of IFN-γ, TNF-α, and IL-2 as biomarkers in a small volume (3µL) of IGRA plasma supernatant samples is demonstrated showing ability to distinguish IGRA+ and IGRA-individuals from a TB endemic setting using this platform.

## MATERIALS AND METHODS

### Materials

Deionized water (DIW, LC267405), phosphate buffer saline (PBS, 21-040-CVR), and reagent alcohol (denatured alcohol, 70% v/v) (25-467-01SC) were obtained from Fisher Scientific. Tween 20 (97062-332), Alconox powder detergent (21835-032), and disposable polystyrene sterile petri dishes (25384-088) were obtained from VWR. Bovine serum albumin (A3294-50G) and biotinyl tyramide (SML2135-50MG) were obtained from Sigma-Aldrich. 1-Step Ultra TMB-ELISA Substrate Solution (34029), Poly-HRP Streptavidin (N200), HRP-Conjugated Streptavidin (N100), RPMI-1640 Medium (A1049101), Fetal Bovine Serum (A5670401), Penicillin-Streptomycin (15140122), and Phytohemagglutinin-L (PHA-L) Solution (00-4977-93) were obtained from Thermo Fisher Scientific. Biotinylated Bovine Serum Albumin (29130) was obtained from Thermo Scientific. Recombinant human IFN-γ protein (11725-HNAS), biotinylated anti-human IFN-γ antibody (11725-R209-B), anti-human IFN-γ antibody (11725-R238), HRP-conjugated anti-human IFN-γ antibody (11725-R209-H), recombinant human TNF-α protein (10602-HNAE), biotinylated anti-human TNF-α antibody (10602-MM08-B), and anti-human TNF-α antibody (10602-MM01) were obtained from SinoBiological. Human IL-2 ELISA Set without plates (ab48471) was obtained from Abcam. Human Peripheral Blood Mononuclear Cells (PBMCs, 70025) were obtained from Stem-cell Technologies. Polydimethylsiloxane film (PDMS, 303718) was obtained from Greene Rubber. EnzMet for General Research Applications (6010-45ML) was obtained from Cedarlane. 96-well microplates (CLS9018) were obtained from Millipore Sigma.

### Clinical Samples

Blood samples were collected from healthy, asymptomatic adults in Kisumu, Kenya for IFN-γ release assay (IGRA) testing using the QuantiFERON-Plus, as previously described^26^. All participants provided written informed consent for participation in the study. Plasma supernatants remaining after IGRA testing were cryopreserved and used to validate our platform.

### PDMS Preparation

A thin PDMS film (0.1 mm thickness) was laser cut to create an array of 2 mm wells, sonicated in 5% Alconox in DIW, then rinsed in DIW. Scotch tape was used to remove any debris from the PDMS before reversible sealing in a petri dish.

### PBMC Stimulation

Cryopreserved PBMCs from US-based healthy donors were commercially obtained (Stemcell Inc), thawed and washed in RPMI-1640 medium supplemented with 10% Fetal Bovine Serum and 1% Penicillin-Streptomycin. 250,000 cells were added to each of 3 wells in a 24-well cell culture plate containing 1 mL media each. 2 µL, 1 µL, and 0 µL of stock PHA-L were added to wells and pipette mixed for PHA-L concentrations of 10 µg/mL, 5 µg/mL, and 0 µg/mL. The 0 µg/mL well served as the unstimulated control. After incubation at 37°C and 5% CO_2_ for 6 days, cell suspensions were removed and transferred to tubes and cells were pelleted. Cell supernatant was pipetted out of the tube without disturbing the pellet.

### Biotin-BSA HRP-SA Model Assay

A serial dilution of biotin-BSA was added to each well in 3 µL drops starting at a concentration of 2.5 µg/mL with 2-fold serial dilutions until 0.076 ng/mL at RT for 2 hours. Next, the slide was blocked with 1% BSA in 0.1% Tween 20 in PBS (0.1% PBST) for 30 min. Blocking buffer was removed and followed by 4 washes with wash buffer (0.05% PBST) by filling the petri dish, swirling by hand for 10-15 seconds, and removing the wash buffer. Poly-HRP Streptavidin diluted to 1 µg/mL in dilution buffer (DB, 0.1% BSA in 0.05% PBST) was added to each well in 3 µL drops and incubated for 1 hour at RT. The petri dish was washed and washed again five times with DIW.

### Cytokine Immunoassay

Cytokine capture antibodies were prepared at 17 µg/mL in PBS (IL-2 capture antibody was prepared at a 1:100 dilution in PBS), added in 3 µL drops to each well in the petri dish, and incubated overnight at 4C. Next, the petri dish was washed three times with wash buffer (0.05% PBST) by filling the dish, swirling by hand for 10-15 seconds, and removing the wash buffer. The dish was blocked with 1% BSA in 0.05% PBST and incubated at RT for 1 hour at 55 rpm on a plate shaker. The petri dish was washed four times in wash buffer as above, and dried. For sample, either serial dilutions of recombinant cytokine protein, stimulated PBMC supernatant, or clinical supernatant sample in DB were added at 3 µL/well and incubated at RT for 2 hrs. The petri dish was washed four times in wash buffer for 1 minute at 85 rpm on a plate shaker, and dried. Biotinylated antibodies at 0.425 µg/mL in DB (IL-2 biotinylated antibody was prepared at a 1:100 dilution in DB) were added at 3 µL/well and incubated for 1 hr. Following washing and drying, poly-HRP Streptavidin at 1 µg/mL in DB was added to each well, incubated for 15 min at RT. Biotinyl tyramide buffer (BTB) was prepared with a 470:30:500 ratio of PBS, Hydrogen Peroxide, and DI water. Following washing and drying, biotinyl tyramide at 0.16 µg/mL in BTB was added to each well, incubated for 10 min at RT. Following washing and drying, poly-HRP Streptavidin at 1 µg/mL was again added to each well, incubated for 15 min. The petri dish was washed and washed again five times with DI water. Traditional plate-based ELISAs were also performed for platform comparison. ELISA protocols are detailed in the Supplementary document.

### Silver Metallization

2 µL drops of enzymatic metallization substrate components A, B, and C were sequentially added and incubated for 4, 4, and 20 min, respectively. To stop the reaction, the petri dish was rinsed with DIW and dried.

## RESULTS AND DISCUSSION

### Assay Platform and Signal Enhancement Cascade Development

Using the known high-binding capacity of polystyrene commonly used for ELISA assays and cell culture, we leveraged this characteristic here by using a commonly available polystyrene petri dish as the substrate for the eMO-BEAM platform. To create individual wells for a high-throughput assay on a petri dish, a custom microwell array was created by laser-cutting 2 mm circles in a thin PDMS film (0.1mm) which is obtained commercially in a roll-based format. This film was reversibly sealed to the inside of a petri dish. The shallow PDMS wells and dish format enable easy wash steps as the dish can be filled with wash buffer and subsequently aspirated or drained. Here, we used a 17 by 5 array of wells for a total of 85 wells, but the PDMS film can easily be cut to enable higher or lower numbers and sizes of wells depending on user need. The microwells are arrayed to make multichannel pipetting possible. This platform enables low sample use (∼3µL) in a low-cost platform which is easy to manufacture using widely available rapid prototyping tools and without the need for any microfabrication facilities.

To establish the feasibility of eMO-BEAM for biomarker detection using enzymatic silver metallization, a model biotin-streptavidin binding assay was performed. Biotin-conjugated bovine serum albumin (biotin-BSA) was incubated on the polystyrene well surfaces as a target molecule and poly-horseradish peroxidase-conjugated streptavidin (poly-HRP-SA) was used as a probe (Figure 1a) followed by enzymatic silver metallization (see Materials and Methods for details). An image of the dried petri dish was captured using a cellphone camera, and the silver darkness was quantified by measuring the grayscale value of the spots formed in ImageJ. A sample dilution dependent silver darkness curve was extracted from plotted grayscale values (Figure 1b-c). Darker spots of silver were found to corresponded with higher biotin-BSA concentrations in a dilution-dependent manner.

**Figure 1.**
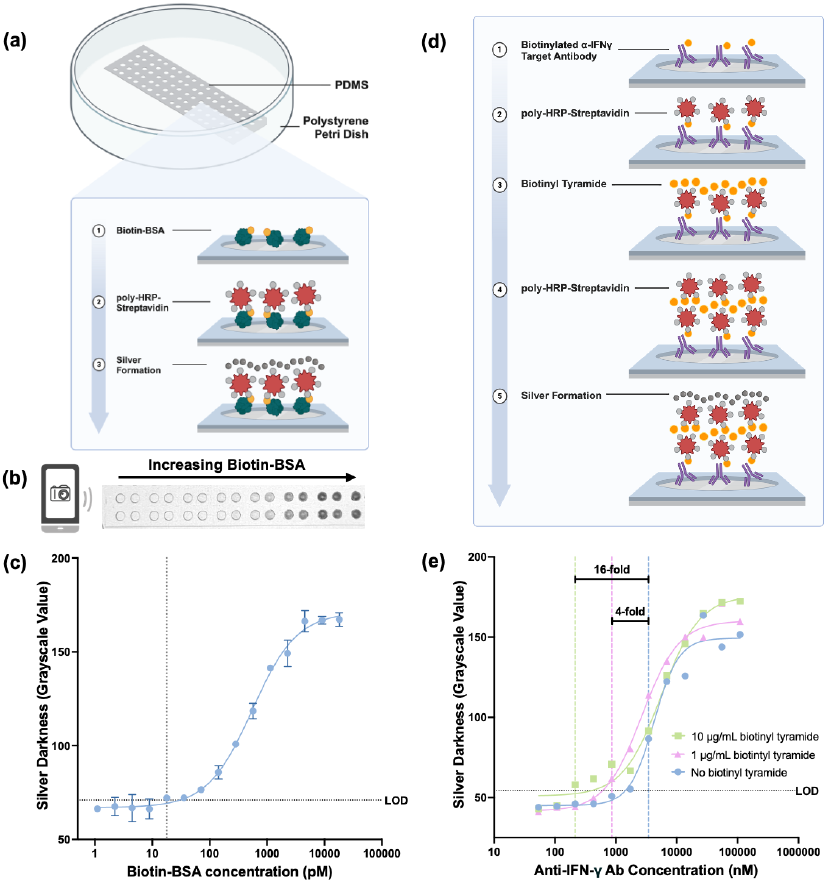
(a) Schematic of laser-cut PDMS microwell array reversibly assembled on a polystyrene petri dish and workflow of enzymatically amplified silver metallization using Biotin-BSA and poly-HRP-SA. (b) Cell-phone image of silver metallized wells after assay. (c) Biotin-BSA dilution curve obtained from cell-phone image silver darkness quantification in ImageJ. (d) Assay stack for initial biotinyl tyramide signal amplification testing using biotinylated anti-IFN-γ as capture. (e) Dilution curve of silver darkness of increasing biotinylated anti-IFN-γ as target and 10 µg/mL, 1 µg/mL, and 0 µg/mL biotinyl tyramide. All assays were repeated three (n=3) times and fit lines are four-parameter logistic sigmoidal curves.

Next, we investigated a signal amplification technique using biotinyl tyramide with serial dilutions of biotinylated anti-IFN-γ antibody as a target molecule in the microwells (Figure 1d). Briefly, after target immobilization, the microwells were incubated with poly-HRP-SA. After a wash, microwells were incubated with biotinyl tyramide to deposit additional biotin sites near immobilized poly-HRP-SA. Following another wash, microwells were again incubated with poly-HRP-SA, and enzymatic silver metallization steps were performed. Comparing a concentration of 10µg/mL and 1µg/mL biotinyl tyramide to the standard method without biotinyl tyramide, we observed a 16-fold and 4-fold improved (i.e. lower) limit of detection respectively confirming this as a viable signal amplification technique for our platform and assay (Figure 1e).

### Cytokine Assay Development

While the above model assays used a biotinylated capture molecule directly immobilized on the well surfaces, next we moved to the full sandwich immuno-assay for cytokine detection (Figure 2a). Initially, an assay without any signal enhancement, using only poly-HRP-SA as a probe was performed with recombinant IFN-γ as the analyte. Dark metallization at the highest IFN-γ concentration, with diminishing signal in serial IFN-γ dilutions, was observed. This resulted in a LOD of 5600 pg/mL being measured (shown in orange in Figure 2b). To amplify the signal and lower the LOD, biotinyl tyramide amplification steps, as above, were added to the assay, resulting in a 127-fold improvement in LOD to 44 pg/mL (shown in blue in Figure 2b). After establishing this cytokine assay protocol with IFN-γ, immunoassays for recombinant IL-2 and TNF-α were performed as well. The LODs for IFN-γ, IL-2, and TNF-α were thus found to be 27.7 pg/mL, 35 pg/mL, and 16.5 pg/mL, respectively which corresponds to 0.71 pM, 2.32 pM and 0.32 pM (Fig 2c-e).

**Figure 2.**
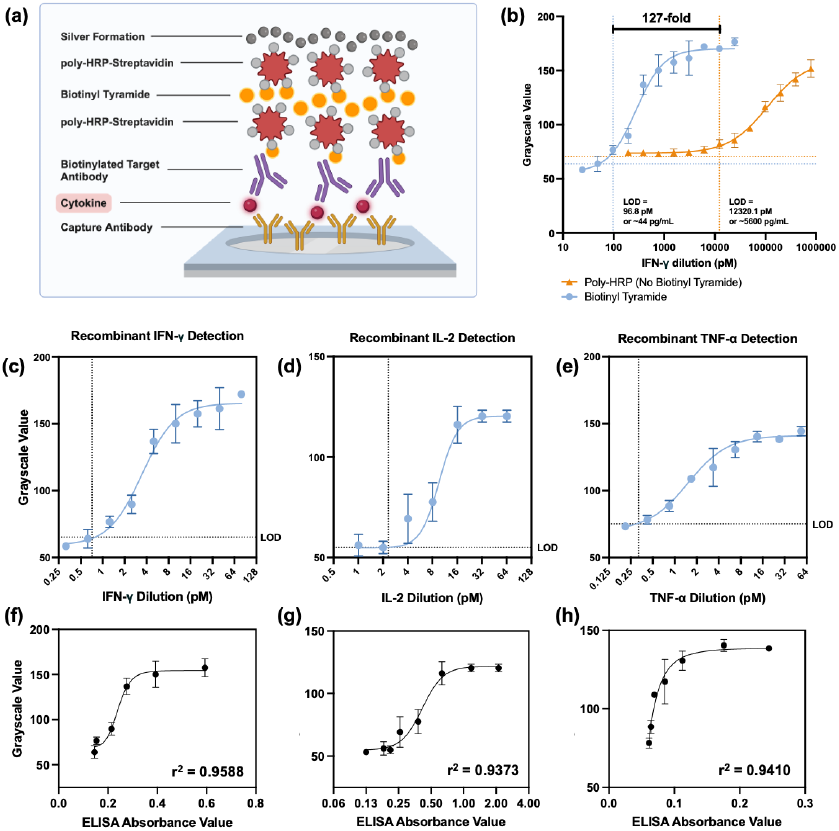
(a) Schematic of immunoassay for cytokine detection. (b) Initial LOD optimization for the IFN-γ assay. (c-e) IFN-γ (n=2), IL-2 (n=5), TNF-α (n=2) dilution curves using recombinant cytokines (f-h) Correlation plots of ELISA absorbance values at 450 nm versus silver darkness values for IFN-γ (f), TNF-α (g), and IL-2 (h). All fit lines are four-parameter logistic sigmoidal curves.

Standard plate-based ELISAs were also performed for these cytokines (Supplementary Figure S1). Correlation plots of grayscale values from our platform against ELISA absorbances generally show a sigmoidal relationship, especially with IFN-γ and IL-2 (Figure 2f-h). At low concentrations, we observe little change in both absorbance and grayscale values, indicating similar LODs between platforms, with ELISA performing slightly better with IL-2 and eMO-BEAM performing slightly better with TNF-α. At higher concentrations, differences are detectable with ELISA, however, silver metallization in eMO-BEAM is observed to saturate, indicating a lower dynamic range compared to ELISA. However, detecting low cytokine levels is more clinically relevant, and sufficient from a diagnostic perspective. Overall, these results establish the enhanced silver metallization on a polystyrene petri dish in eMO-BEAM as a viable platform for cytokine detection, even at low concentrations.

### Cytokine Detection in Stimulated Primary Human Immune Cell Supernatant

To validate the use of this platform with primary human immune cells, cytokine detection in supernatant extracted from stimulated peripheral blood mononuclear cells (PBMCs) from healthy donors was performed (see Materials and Methods for details). Briefly, PBMCs were incubated with phytohemagglutinin-L (PHA-L), a mitogen that binds to immune cell surface receptors and stimulates cytokine production, in a cell culture plate (Figure 3a). Cell culture supernatant from these cells was then used as a sample on our platform, at a range of sample dilutions, for detection of IFN-γ, TNF-α, and IL-2. The stimulated and unstimulated cells showed a clear difference in the measured signal (i.e. silver darkness) which reduced in a sample dilution-dependent manner and was detectable in sample dilution factors of up to 128 (Figure 3b-d). Additionally, an effect of mitogen concentration was observable as well. Interestingly, we observed a saturated signal in TNF-α detection down until a sample dilution factor of 128 using 10 µg/mL PHA-L and 64 using 5 µg/mL PHA-L, indicating a high level of TNF-α produced by stimulated PBMCs. On the other hand, IL-2 was only detectable in the 10 µg/mL PHA-L stimulated sample, and the signal only begins to saturate at the undiluted sample concentration, suggesting this mitogen may not stimulate large amounts of IL-2 secretion at lower mitogen concentrations. Additionally, at high sample concentrations we observed that IFN-γ was detectable in the samples from unstimulated cells as well which can represent the baseline level of this cytokine secreted by these cells. Ultimately, these results indicate that the eMO-BEAM platform can detect natural primary human immune cell-secreted cytokine with minimal to no cross-reactivity from any other substances that may be present in cellular supernatant, as validated by lower signal from unstimulated samples.

**Figure 3.**
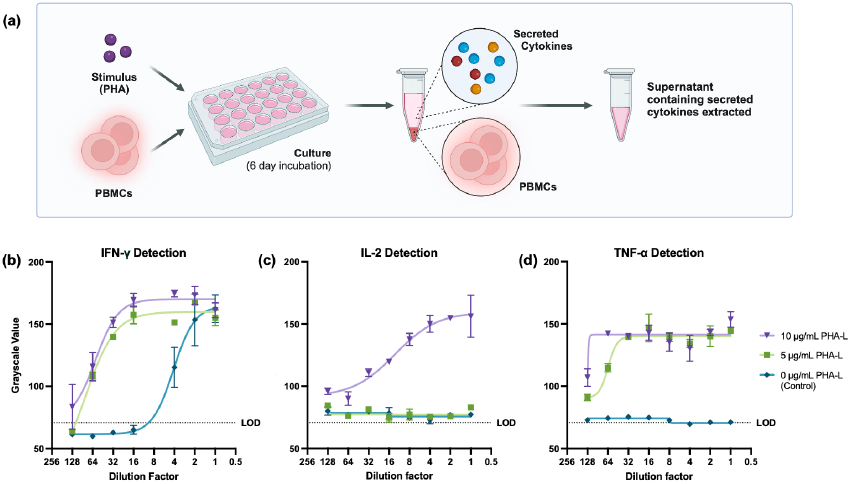
(a) PBMC stimulation workflow using PHA-L as mitogen and a 6-day incubation. Cells were either incubated with 10 µg/mL, 5 µg/mL, or 0 µg/mL (control) PHA-L. Cells were pelleted and supernatant containing cytokines was extracted. (b-d) Dilution curves for cytokine detection in cell supernatant (n=2). All fit lines are four-parameter logistic sigmoidal curves.

### Cytokine Detection in Clinical Samples

Next, to test the performance of the eMO-BEAM platform with clinical samples, we measured the cytokine levels in a set of IGRA supernatant samples from adults in Kenya (IGRA+, n=3 and IGRA-, n=4). A group of four stimulated supernatant samples was available for each participant: NIL (unstimulated negative control), Ag1, Ag2, and Mitogen (positive control), as these are the set of peptide preparations used for stimulation in the IGRA test performed for these participants (Figure 4a). Ag1 is known to be the set of Mtb antigen ESAT-6 and CFP-10 peptides targeting CD4+ T-cell response while Ag2 contains additional antigen peptides to also target CD8+ T-cell response^27, 28^. First, serial dilutions of pooled participant samples (i.e. 4 pools each for IGRA+ and IGRA-respectively) were used to measure IFN-γ, TNF-α, and IL-2 (Figure 4b-e). Generally, good differentiation was observed between the IGRA+ and IGRA-samples for the Ag1 and Ag2 stimulated samples for all three cytokines. Overall, these pooled sample serial dilution curves indicated that a sample dilution factor between 10 to 20 can effectively be used to distinguish between stimulation conditions, as well as between IGRA+ and IGRA-samples in this assay. We also observed that mitogen-stimulated samples exhibited relatively high signal compared to NIL, Ag1, and Ag2 samples throughout IFN-γ and TNF-α dilution curves, as expected. However, in the IL-2 dilution curve, mitogen-stimulated sample was similar or lower than some antigen-stimulated samples for the IGRA+ pools; this could suggest that the mitogen used did not stimulate IL-2 secretion in additional amounts in these participant samples (Figures 4d-e). Additionally, considering the NIL or unstimulated controls, relatively higher levels of TNF-α, but not IFN-γ or IL-2 were observed in these samples which could represent higher baseline levels of secretion of this cytokine in these samples.

**Figure 4.**
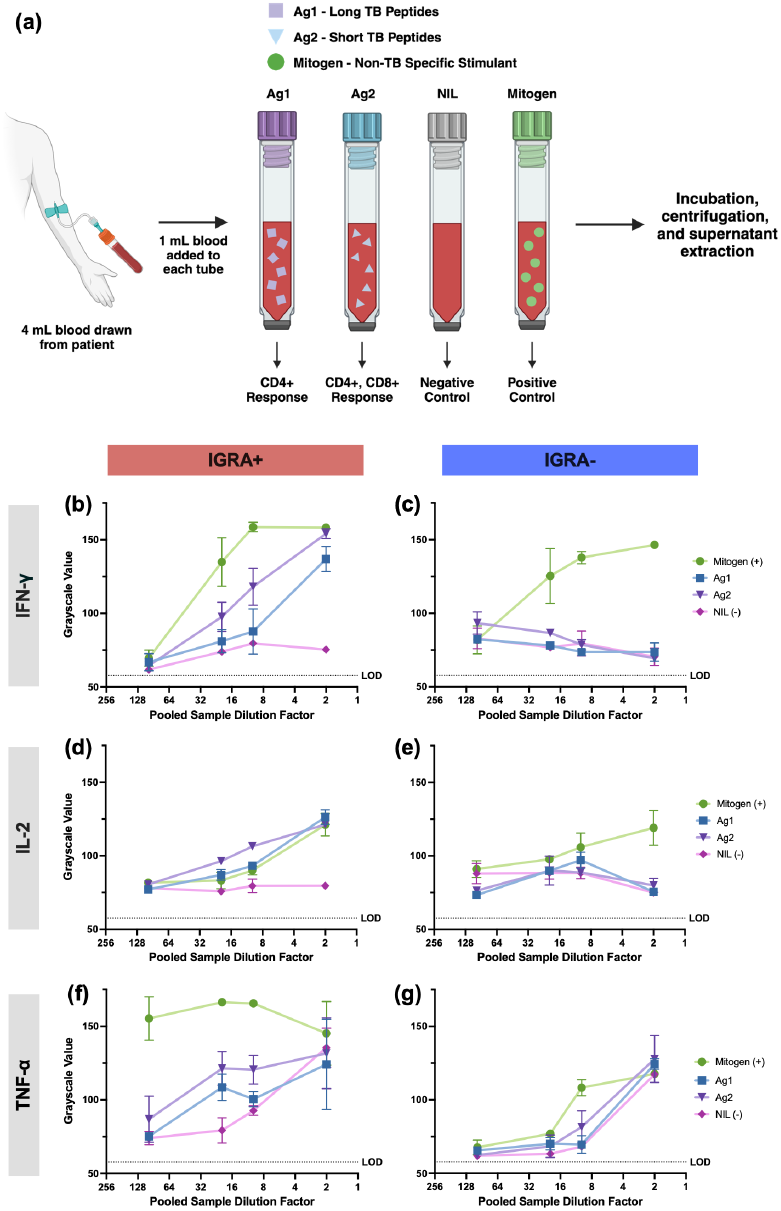
(a) Schematic demonstrating the stimulation groups and controls. Dilution curves for IFN-γ (b-c), IL-2 (d-e), and TNF-α (f-g) in pooled LTB+ and LTB-clinical supernatant samples. Clinical samples consisted of NIL (negative control), Ag1, Ag2, and Mitogen (positive control) groups (n=2).

After the above sample dilution factor optimization using pooled IGRA+ and IGRA-samples, individual participant samples were run on the eMO-BEAM platform (using a sample dilution of 10), and each stimulated supernatant sample group was compared based on participant IGRA status (Figure 5). This revealed several interesting observations. For IFN-γ, the mitogen and NIL samples exhibited high and low silver signals, respectively, as expected. Ag2-stimulated samples showed significantly higher IFN-γ levels in all IGRA+ vs all IGRA-participants. Ag1-stimulated samples performed worse in differentiating between IGRA statuses, with 2 of 3 IGRA+ individuals showing higher IFN-γ levels with Ag1 stimulation. Notably, IL-2 was found here to be the most effective biomarker in differentiating this set of IGRA+ participants from IGRA-participants. IGRA-Ag1 and Ag2 sample IL-2 levels were near NIL levels, while IGRA+ IL-2 signals in Ag1 and Ag2 were significantly high throughout all patients. TNF-α detection in Ag1 and Ag2 samples show significant differences between IGRA+ and IGRA-individuals as well. However, we also observe high TNF-α signal in the IGRA+ NIL (unstimulated) group, suggesting high baseline TNF-α levels in IGRA+ individuals. TNF-α is known to play a key role in containing the spread of bacteria, and therefore with in latently infected individuals, TNF-α is likely constantly present in relatively high amounts to contain the dormant bacteria^29^. Results from a commonly used calculation for IGRA tests, where the NIL values are subtracted from the antigen-stimulated values, are shown in Supplementary Figure S3 as well. Overall, these measurements and the heterogeneity in differences in the three cytokine levels between IGRA+ and IGRA-individuals underscore further the value of multiplexed cytokine detection in clinical diagnostic use.

**Figure 5.**
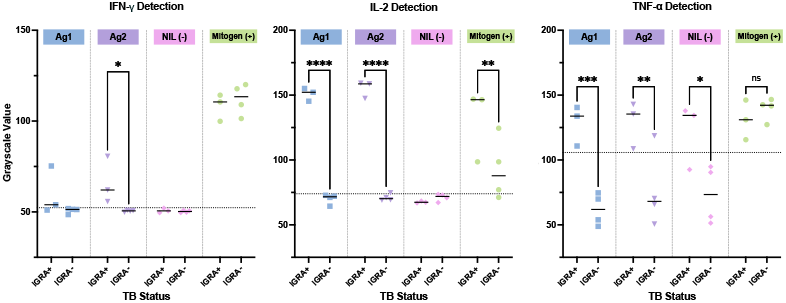
Individual participant silver darkness values for IFN-γ (a), IL-2 (b), and TNF-α (c) detection from NIL, Ag1, Ag2, and Mitogen samples of IGRA+ and IGRA-patients using a sample dilution of 1:10. Average of n=2 repeats was plotted for each sample.

### Multiplexed Cytokine Assay Development and Clinical Sample Testing

Finally, we tested the capability of a multiplexed format of this device to measure all three cytokines from a single drop of sample as shown in Figure 6. To do so, two layers of PDMS were laser-cut; one containing four smaller (diameter: 1.5 mm) microwells, and one containing a larger (diameter: 6 mm) sample microwell. The layer containing the 4 small microwells was reversibly sealed under the petri dish and served as a visual alignment template for capture antibody droplet placement inside the petri dish. The layer containing the large wells was reversibly sealed inside the petri dish and served as a common well for a single drop of sample and all following reagents (Figure 6a). This format was first tested for any potential cross-reactivity using recombinant cytokines of known concentrations. Here four small capture spots were then created using 1.5 µL droplets of anti-IFN-γ, anti-TNF-α, and anti-IL-2 capture antibodies, as well as a 1X PBS buffer droplet to serve as a negative control. Equal volumes of the different recombinant cytokine samples were then added to the larger wells, followed by a common mixture of secondary antibodies, signal enhancement and enzymatic metallization reagents as earlier (Figure 6, see Materials and Methods section). Silver metallization was only observed in the cytokine-specific capture antibody droplet spots. Thus, for example, only the anti-IFN-γ spot was dark in the sample well which received IFN-γ as the sample. Also, the anti-IFN-γ spots were not dark in the other sample wells which received TNF-α and IL-2 as sample. This was true for each of the other cytokines too (Figure 6b). Additionally, none of the control spots were dark and none of the spots in the control well was dark either. This thus confirmed the specificity of assay reagents and minimal cross-reactivity effects and overall established the ability to detect multiple cytokines in the eMO-BEAM platform from a single drop of sample.

**Figure 6.**
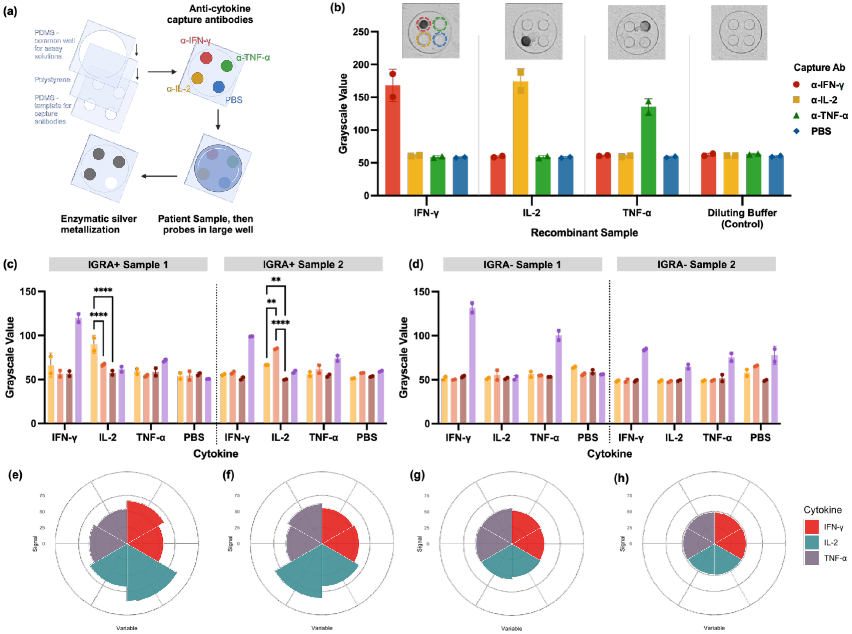
(a) Workflow for multiplexed detection of cytokines with a common well for sample/recombinant protein, probes, and enzymatic reagents. (b) Demonstration of minimal cross-reactivity using recombinant cytokines. X-axis indicates the recombinant protein used as sample in large common well. Only the corresponding capture antibody spot produced silver signal. (c) Multiplexed cytokine testing in individual IGRA+ (n=2) and (d) IGRA-(n=2) samples. (e-h) Multiplexed cytokine fingerprints of individuals.

Using this multiplexed assay format, simultaneous detection of IFN-γ, TNF-α, and IL-2 was then performed from a subset of IGRA supernatant samples. The results of this are shown in Figure 6c and d. Overall, IGRA+ samples were clearly distinguishable from IGRA-samples based on this measured multiplexed cytokine fingerprint as well (Figure 6e-h). Several other interesting observations also were made in this measurement. IL-2 levels again showed good differentiation between TB antigen-stimulated samples and the NIL sample with IGRA+ but not IGRA-individuals. This matches the earlier observation shown above in the single-plex assays (Figure 5b). However, with IFN-γ and TNF-α, we did not see such clear differentiation. With IFN-γ, this matches the earlier single-plex data as well (Figure 5a). However, the TNF-α detection in the multiplexed assay format did not perform as well as the earlier single-plex detection. This could be due to further need for assay optimization here. Since the secondary antibodies for all three cytokines were mixed due to the multiplexed nature of the assay, there may be a more optimal concentration of the anti-TNF-α antibody needed. Alternatively, a more concentrated sample dilution may be worth exploring for better performance in the multiplexed platform. Despite this difference, overall, we were able to validate the use of this platform for multiplexed detection of cytokines from single drops of individual patient samples.

Additionally, overall IL-2 outperformed the other cytokines in both single-plex and multiplexed versions of our platform, allowing for better differentiation between this set of IGRA+ and IGRA-individuals. While only IFN-γ is currently used in commercial LTBI diagnostic assays, this further shows the value of multiplexed measurement of other cytokines.

## Conclusions

We have presented here a low-cost yet highly sensitive (sub-picomolar), multiplexable biosensing platform for low abundance cytokine biomarker detection. Compared to our prior work with antibody detection, MO-BEAM, this represents a 1000-fold increase in sensitivity and thus we term it as **<u>e</u>**nhanced MO-BEAM or eMO-BEAM. The eMO-BEAM platform is developed using a common petri dish and a commercially obtained roll of PDMS thin sheet which are both easily attainable and storable consumables that do not need any specialized microfabrication or other manufacturing techniques except for simple rapid prototyping and general tools widely available globally.

The signal enhancement cascade reported here enables sensitive detection of low abundance biomarkers with the dry silver darkness-based readout which is obtained using a cellphone camera. We have earlier shown that this can be analyzed and quantified using a custom cellphone app we have developed and made available as open source software as well^30^. This represents a significant reduction in instrument as well as consumable cost compared to other ELISA or multiplexed bead-based assay techniques which require optical detection instrumentation such as a plate reader or flow cytometer as well as need larger amounts of expensive reagents like antibodies and barcoded beads. A summary of key feature comparisons between ELISA, multiplexed bead-based assays and the eMO-BEAM platform is presented in Table 1. It is worth noting that while some manual assay steps (e.g. pipetting) are still needed in the current form of this platform, as are needed for other comparable techniques, these can be automated, if needed, in the future using one of several inexpensive, open hardware or cartridge-based solutions others have developed^31-35^.

**Table 1.** Assay comparison between ELISA, Bead-based Assays and eMO-BEAM.

| Method | Multiplexable | Sample Volume (μL) | Consumables Cost | Instrument Cost | Time (hrs) | Sensitivity | Portable |
| --- | --- | --- | --- | --- | --- | --- | --- |
| ELISA | No | 50 | ~\$10 | \$20K | 3-4 hours | pM | No |
| Bead-based Assays | Yes | 50 | ~\$10 | \$30K - \$150K | 3-4 hours | pM | No |
| eMO-BEAM | Yes | 3 | ~\$0.1 | <\$100 | 4-5 hours | pM | Yes |

Furthermore, we applied the eMO-BEAM platform here for detecting low concentrations of multiple cytokine biomarkers (IFN-γ, TNF-α, and IL-2) in stimulated human primary immune cell supernatants from cell culture as well as directly from small volumes (<5µL) of clinical samples, which are compatible with the commonly used IGRA test. By significantly decreasing the amount of lab infrastructure needed to perform such a multiplexed cytokine measurement while increasing the amount of data attainable from one drop of patient sample, this platform creates the opportunity to perform cytokine detection in labs in resource-poor settings. In conclusion, the characterization and performance of this biomarker detection platform demonstrates its diagnostic capability, and its tunable properties hold potential for adaptability for point-of-care detection of other disease biomarkers beyond TB as well.

## ASSOCIATED CONTENT

## AUTHOR INFORMATION

### Author Contributions

The manuscript was written through contributions of all authors. All authors have given approval to the final version of the manuscript. ‡These authors contributed equally to this work as co-first authors and retain the right to list themselves first in author order on their CVs.

