## Supplementary Information for "Inexpensive High-Throughput Multiplexed Cytokine Detection for Tuberculosis Diagnostics Using Amplified Enzymatic Metallization"

#### Supplementary Methods

All ELISA kits and matched antibody sets are mentioned in Materials & Methods.

**IFN-γ ELISA:** 96-well plates were coated overnight at 4C with 50 µL/well of 2 µg/mL anti-IFN-γ capture antibody in PBS. The plates were washed three times with 0.05% PBST and blocked with 1% BSA in 0.05% PBST for 1 hour at room temperature (RT) followed by three washes with 0.05% PBST. Serial dilutions of recombinant IFN-γ protein diluted in 0.05% PBST were added at 50 µL/well. After 2 hrs of incubation at RT, wells were washed three times with 0.05% PBST. __ µg/mL of anti-IFN-γ-HRP was added at 50 µL/well and incubated for 1 hr at RT. Following three washes with 0.05% PBST, 50 µL of TMB substrate was added. After 15 minutes, the reaction was stopped by adding 50 µL of 1 M sulfuric acid. Absorbance was read at 450 nm using a microplate reader.

**TNF-α ELISA:** 96-well plates were coated overnight at 4C with 50 µL/well of 2 µg/mL anti-TNF-α capture antibody in PBS. The plates were washed three times with 0.05% PBST and blocked with 1% BSA in 0.05% PBST for 1 hour at room temperature (RT) followed by three washes with 0.05% PBST. Serial dilutions of recombinant TNF-α protein diluted in 0.05% PBST were added at 50 µL/well. After 2 hrs of incubation at RT, wells were washed three times with 0.05% PBST. 0.5 µg/mL of anti-TNF-α-Biotin was added at 50 µL/well and incubated for 1 hr at RT. After 1 hr of incubation at RT, wells were washed three times with 0.05% PBST. 0.007 µg/mL of HRP-Streptavidin was added at 50 µL/well and incubated for 1 hr at RT Following three washes with 0.05% PBST, 200 µL of TMB substrate was added. After 15 minutes, the reaction was stopped by adding 50 µL of 1 M sulfuric acid. Absorbance was read at 450 nm using a microplate reader.

**IL-2 ELISA:** IL-2 ELISA was performed according to manufacturer protocol.

### Supplementary Figures

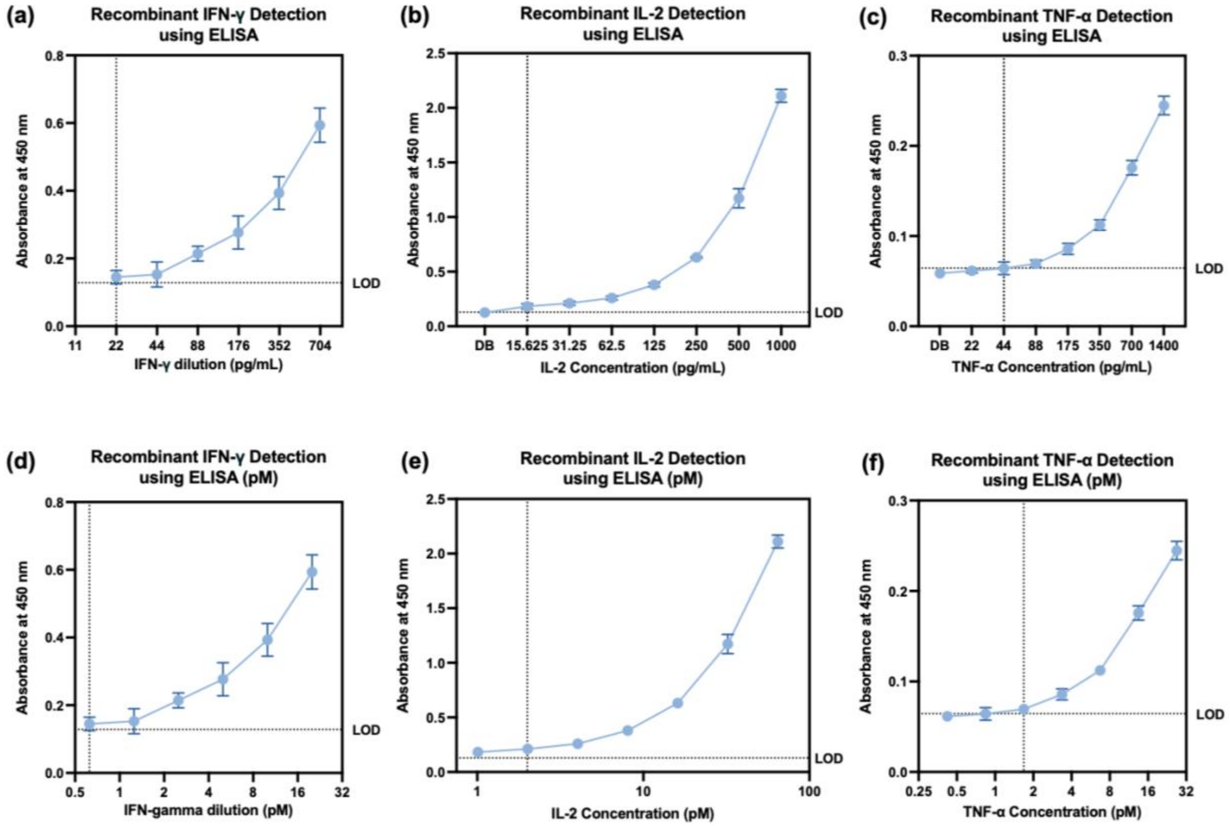


**Figure S1.** ELISA for recombinant cytokine detection and LOD derivation. LODs of 22, 15.625, and 44 pg/mL were determined for IFN-γ (a), IL-2 (b), and TNF-α (c), respectively. pM converted graphs show LODs of 0.625, 2.016, and 1.683 pM for IFN-γ (d), IL-2 (e), and TNF-α (f), respectively.
